# Testing the fast consolidation hypothesis of retrieval-mediated learning

**DOI:** 10.1101/458687

**Authors:** C.S. Ferreira, I. Charest, M. Wimber

## Abstract

The testing-effect, or retrieval-mediated learning, is one of the most robust effects in memory research. It shows that actively and repeatedly retrieving information, compared to merely restudying it, improves long-term retention. Surprisingly, little is known about the neurocognitive mechanisms underlying this phenomenon. Attempting to fill this gap, a recent framework proposed that retrieval acts as a fast memory consolidation mechanism, stabilizing memories through online reactivation, similar to memory replay during offline (e.g. sleep) consolidation. In this fMRI study, we empirically tested the predictions derived from this framework.

We predicted that reactivation during retrieval allows memories to become embedded in neocortex, creating an additional route to access the memory trace and rendering it less hippocampus-dependent. Participants encoded scene-object pairs and either retrieved or restudied the objects over two sessions, two days apart. We analysed univariate and multivariate changes in brain activity specific to retrieval but not restudy, and tested whether the predicted changes occur rapidly within a session, or evolve slowly, across the two days.

Results showed that medial prefrontal cortex activation increased across retrieval trials within one session, consistent with a fast consolidation account. Hippocampal activity decreased across sessions, suggesting a slower mechanism. Moreover, Representational Similarity Analyses (RSA) showed that consecutive retrieval attempts strengthen both higher-level semantic and episode-specific information in parietal areas, both across but not within sessions. Our findings suggest that retrieval supports the online creation of a neocortical trace, which becomes increasingly relevant at long delays when hippocampus-dependent episodic details would otherwise have faded.

**Significance statement:** Repeated remembering strengthens memories much more so than repeated learning. The aim of this study was to shed light onto the poorly understood neurocognitive underpinnings of retrieval-mediated learning. We tested a novel framework proposing that a memory’s stabilization via retrieval relies on mechanisms akin to those involved in offline systems consolidation. Observing the retrieval-induced neural pattern changes across different timescales, we find that retrieval stabilizes semantic and episodic aspects of the original memories, and produces increases in prefrontal activity and decreases in hippocampal activity that are consistent with the consolidation view, but not necessarily with a fast acting mechanism. Our findings inform cognitive theories of the testing effect, suggesting that retrieval produces its benefits by interacting with hippocampal-neocortical consolidation mechanisms.

## Introduction

Actively and repeatedly retrieving a newly acquired memory is a powerful tool to promote its long-term retention, compared to merely restudying it. This retrieval benefit, termed the testing-effect or retrieval-mediated learning, has been replicated in countless studies, using different types of materials, learners and contexts (Roediger & Karpicke, 2006; Karpicke, Lafayette, & States, 2017).

Surprisingly, few cognitive theories offer a mechanistic explanation for retrieval-mediated learning (Karpicke et al., 2017), and as of yet, none has gathered robust support. Attempting to bridge this gap, we recently proposed that retrieval’s benefits rely on online reactivation mechanisms similar to those involved in offline systems consolidation (Antony, Ferreira, Norman, & Wimber, 2017).

Systems-level consolidation theories propose that the hippocampus and neocortex act synergistically during offline periods (such as sleep) to stabilize mnemonic representations (McClelland, McNaughton, & O’Reilly, 1995). During offline periods, memories are reactivated or replayed in hippocampal-neocortical networks. Reactivation allows new memories to get embedded into pre-existing cortical knowledge structures, creating a neocortical trace more durable than the initial hippocampal one (Dudai, Karni, & Born, 2015). Thus, whereas recent memories rely mostly on the hippocampal trace, remote memories can be accessed via this additional neocortical route, becoming less hippocampus-dependent.

Retrieval and offline consolidation share remarkable theoretical and computational properties, and produce similar behavioural benefits. This led to our proposal that retrieval might act as a fast online consolidation mechanism, and that the creation of a stable neocortical representation is supported by the neural reactivation of memories, rather than by time or sleep *per se* (Antony et al., 2017). This account offers a neurobiologically plausible mechanism for retrieval-based strengthening, but has yet to be tested empirically. In this fMRI study, participants performed consecutive retrieval and restudy blocks over two sessions, two days apart, designed to test three predictions derived from the fast consolidation framework.

First, if an additional cortical trace is created during retrieval, neocortical activation should increase across cycles in the medial prefrontal cortex (mPFC), mirroring sleep-dependent consolidation results (Nieuwenhuis & Takashima, 2011). If memories can then be accessed via this neocortical route, the hippocampus should be required less and its activation should decrease across retrieval attempts.

Secondly, if retrieval embeds memories in neocortex, retrieved items should become more generalised or gist-like (Rasch et al., 2007; Richards et al., 2014; Schapiro et al., 2017). Empirically, the neural patterns of memories sharing semantic information (items belonging to the same semantic category) should thus become increasingly similar across repeated retrievals.

Finally, given retrieval’s role in reducing neural overlap between similar memories (Antony et al., 2017; Hulbert & Norman, 2015), the fast consolidation account predicts that retrieval strengthens episode-unique information. This should be reflected in increased similarity between an individual item’s neural pattern at study and its reactivated patterns across retrievals, compared with other overlapping memories. Evidence regarding this prediction is conflicting in the systems consolidation literature. While some studies found that, as memories become gist-like (i.e. consolidated), they lose contextual detail (Cairney, Durrant, Musgrove, & Lewis, 2011), others found no evidence for such decontextualization (Jurewicz, Cordi, Staudigl, & Rasch, 2016). *Semanticization* might thus not necessarily lead to a concurrent loss of mnemonic detail. Semantic and episodic traces might instead co-exist.

For each prediction, we were specifically interested in the timescale at which neural patterns evolve. If retrieval acts as a fast online consolidation event, neural changes should occur across repeated retrievals within the first experimental session. Alternatively, if these changes happen in a slower fashion (across days), this would suggest that retrieval’s benefits depend on its interaction with offline consolidation mechanisms.

This is the first study specifically testing the fast consolidation account of retrieval-mediated learning, by measuring neural changes across fast (within-session) and slow (between-days) timescales. Understanding whether retrieval benefits memory through a fast online mechanism, or rather through the interaction with slower consolidation processes, constitutes an important step in unravelling the neurocognitive mechanisms underlying retrieval-mediated learning, offering a neurobiological basis for existing testing-effect accounts.

## Methods

### Participants

Twenty-four volunteers (17 female, M_age_ = 23.3, SD = 4.0) completed the two sessions of the experiment. Two participants were excluded, one due to extreme movement (more than twice the voxel size) and the other for reporting falling asleep in the scanner. Due to technical errors, the third practice cycle of the experiment is missing in one participant, and the fourth cycle in another. The remaining data from these two participants was fully included in the analyses. All 22 participants in the final sample were right-handed, native or very fluent English speakers, had normal or corrected-to-normal vision and no history of neurological, psychological or psychiatric conditions. Participants received course credit or a monetary reward for taking part in the experiment. The experiment was approved by the STEM Ethics Committee of the University of Birmingham.

### Materials

A set of 128 scenes and 128 objects were chosen as stimuli. Each object and scene was unique, but objects belonged to eight different semantic categories: animals, musical instruments, fruits, clothes, sports gear, office supplies, kitchen appliances and furniture, with 16 exemplars per category. The objects were chosen from the Bank of Standardized Stimuli (BOSS; Brodeur, Guérard, & Bouras, 2014; https://sites.google.com/site/bosstimuli/), resized to 170×170 pixels and modified to greyscale. The scenes were drawn from the SUN database (Xiao, Hays, Ehinger, Oliva, & Torralba, 2010; http://groups.csail.mit.edu/vision/SUN/). All scenes were resized to 256×256 pixels and displayed in greyscale. Stimuli were presented using in-house Python code, running on PsychoPy v.1.84.2 (Peirce, 2006; http://www.psychopy.org/).

### Procedure

Participants were asked to complete two sessions on two different days (Figure 1A). On the first session, participants were provided with task instructions and MRI information and asked for their informed consent.

**Figure 1:**
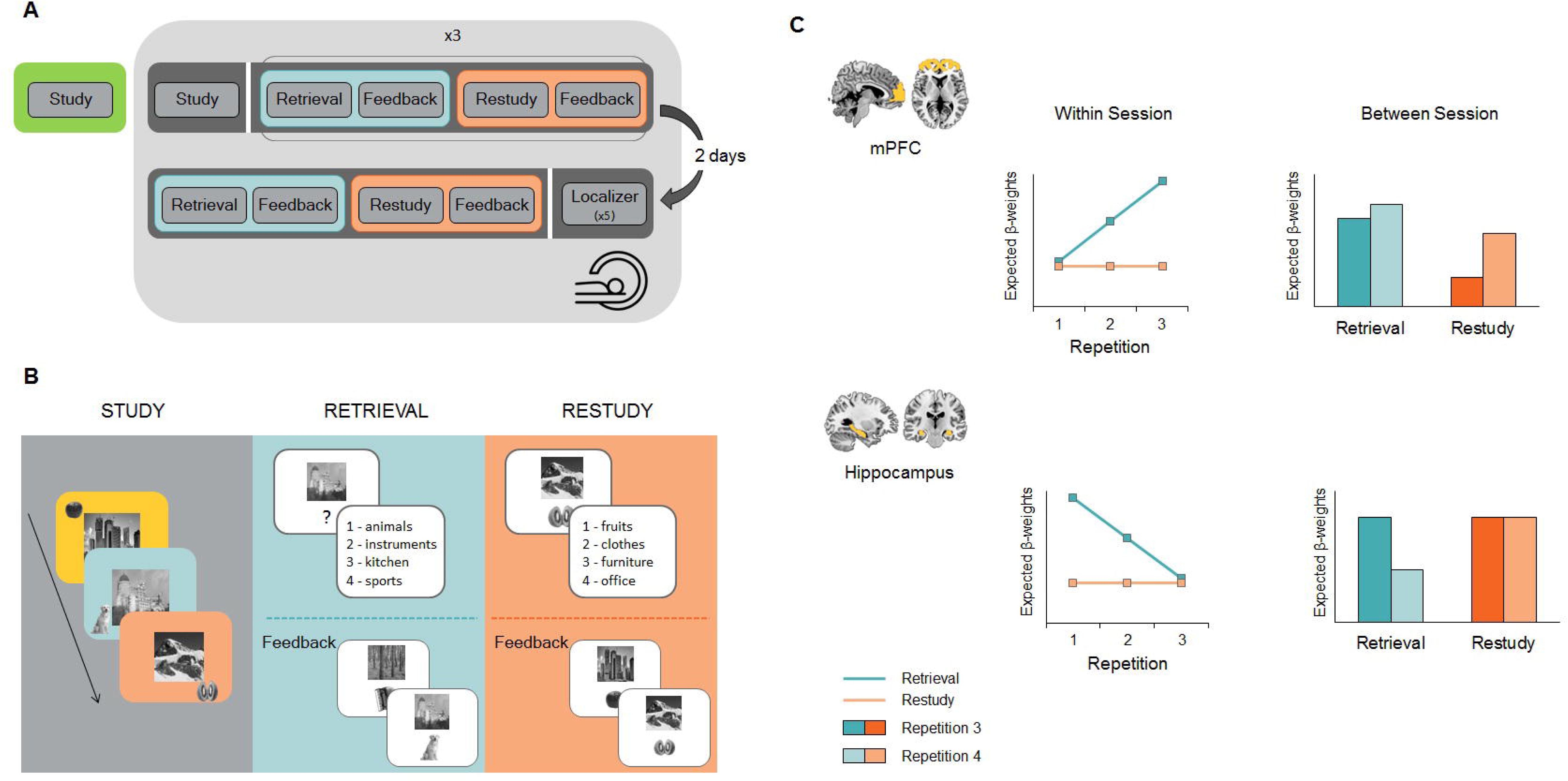
Experimental Procedure. **A.** General timeline of the experiment. Participants were tested on two different days, two days apart. In the first session, participants performed a study phase outside the scanner and another inside the scanner, followed by three cycles of retrieval and restudy trials. Each retrieval/restudy cycle was completed by a massed feedback phase. In the second session, participants performed an additional retrieval and restudy cycle followed by a functional localizer. All the phases encompassed by the light grey background took place inside the MRI scanner. **B.** Detailed depiction of the main phases of the experiment. During the study phase, participants saw different object-scene pairs that they were asked to mentally link together as vividly as possible. The scenes and objects were unique but, importantly, the objects belonged to a number of different semantic categories. During a retrieval trial participants saw a scene and were asked to think back, as vividly as possible, to the object associated with that scene, and to then indicate its semantic category by a button press. Restudy trials unfolded in a similar way, the only difference being that the scene-object pair was presented intact on the screen and thus there was no need for participants to search their memory for a target object. A massed feedback phase followed each cycle, where subjects saw all the intact pairs again, presented in the centre of the screen. **C.** Prediction for univariate effects in each ROI, separately for fast within (left) and slow between (right) session changes. Predicted results for retrieval are depicted in blue and for restudy in orange. Third repetition trials are presented in a darker hue than fourth repetition trials. The top panel shows predictions for mPFC activation, the bottom one for hipopcampus.

In Session 1, participants performed a study phase twice: once outside and once inside the scanner. This was followed by three blocks of practice, each comprising one retrieval and one restudy cycle on separate sets of the studied material. Participants then returned after two days (approximately 48h) for Session 2. In this session they completed an additional practice block (one retrieval cycle and one restudy) and a functional localizer, all inside the scanner.

#### Session 1

##### Familiarisation Phase

Before starting the actual experiment (i.e., the study and practice phases), participants were given an opportunity to get familiar with the tasks. The trials during familiarisation followed the exact same structure as the ones presented later during the experiment, but used 10 stimuli only (5 scene-object pairings) that were used specifically for this phase and never again seen during the remainder of the experiment. None of the objects belonged to any of the semantic categories presented in the experiment itself.

##### Study Phase

At the beginning of the study phase, participants were informed that they would see a series of scene-object pairs that they needed to commit to memory the best they could. Participants were instructed that, to achieve this, they should link the two (scene and object) together as vividly as possible by, for instance, mentally integrating the object into the scene. They were explicitly told that although each object and scene were unique, the objects belonged to a number of different semantic categories. After a 5-trial practice (familiarisation phase), subjects performed the first study phase for all 128 scene-object pairs in a quiet room in our imaging facilities. Participants sat in front of a laptop to perform the task.

A study trial began with a black fixation cross on a white background (jittered 0.5–7.5 sec). This was followed by a scene-object pair, presented in random order, for 4.5 sec. The scene was always shown in the centre of the screen, while the object was presented in one of the four corners. The pair was shown on a different coloured background – pink, blue, green or yellow (Figure 1B). The position of each object and the colour of the background were pseudo-randomly assigned, so that each colour and position would be equally distributed across categories and later practice conditions. Spatial positions and background colours were used to create a more unique encoding context for each item. For each pair, participants were asked to press a key on the keyboard to indicate whether or not they had been able to come up with a vivid mental image connecting the scene and the object.

After the first study phase, participants were taken to the scanner where, after the acquisition of the structural images (∼5 minutes), they performed the second study phase for all object-scene pairings, this time inside the scanner. Images were projected on a screen behind the scanner bore that participants saw through a mirror attached to the head coil. This phase followed the exact same procedure as the one outside the scanner, but stimuli were presented in a different random order. That is, the scene-object pairing, the object position and the background colours were kept constant, but the order of stimulus presentation was changed. Inside the scanner, responses were made on a button box that participants held in their right hand.

##### Practice Phase

The Practice Phase immediately followed Study. In Session 1, this phase comprised three retrieval and three restudy cycles. The order of conditions was counterbalanced across subjects, with half of the subjects performing retrieval-restudy, restudy-retrieval, retrieval-restudy, and the other half following the opposite scheme. Assignment of stimuli to conditions was also counterbalanced across subjects. Each subject performed one of four possible counterbalancing schemes. In each scheme, half of the stimuli (64), belonging to 4 semantic categories, were attributed to retrieval. The other 4 categories (64 stimuli) were assigned to restudy. Stimuli were allocated to retrieval or restudy pseudo-randomly so that each semantic category (with its 16 examplars) was assigned to the retrieval condition in two of the counterbalancing schemes, and to the restudy condition in the remaining two schemes. Each given scene-object pair remained in either the retrieval or restudy condition across all four retrieval or restudy repetitions, across the two days. The order of stimulus presentation was randomized within participant for each cycle.

A retrieval trial (Figure 1B) started with a black fixation cross in the centre of a white screen (jittered 0.5–7.5 sec) and was followed by the presentation of the scene, as a cue to retrieve the object. The scene was presented over a white background, centred in the upper part of the screen with a black question mark below. Participants were instructed to think back to the item associated with this particular scene, and visualize it as vividly as possible in their mind for the full duration of the trial. Afterwards, participats saw four possible categories written on the screen (black font over white background) and were asked to indicate the category to which the object associated with the scene they had just seen belonged to. For a given participant, the categories shown at retrieval and restudy, as well as the button associated to the response, were kept constant throughout the experiment. After the 64 trials of a retrieval cycle, there was a massed feedback phase, where participants saw all the pairs together again, presented for 2 sec each. The scene was presented on the upper part of the screen, with the object centered below it, both over a white background. Participants were asked to press a key during these two seconds to indicate whether the object corresponded to the one they had thought of earlier or not. Massed feedback was included since it has been shown to enhance the retrieval-mediated learning effect (Roediger & Karpicke, 2006). Moreover, this manipulation allowed us to collect an additional (subjective) measure of participants’ performance.

Restudy trials were very similar to retrieval ones. The only difference was that instead of the scene being paired with a question mark, participants saw the whole pair intact again (the scene in the upper part of the screen, with its corresponding object below, over a white background). Participants were instructed to take these trials as an opportunity to relearn the pairs. When the four categories appeared after each trial, they were asked to chose the category that the object of the current trial belonged to. After 64 restudy trials there was also a “feedback” phase, where participants saw all the restudy pairs again. In this case, they were asked to press a button to indicate whether they still found it easy or hard to link the object and scene together. The trial structure of the restudy condition was thus highly similar to the retrieval condition, but involved no active retrieval demand at any point.

After each block of retrieval + restudy, there was a 2 minute break where participants were told to close their eyes if needed and rest. The first session ended after three cycles of retrieval and three of restudy practice. Participants were taken out of the scanner and sent home, with a reminder of the second session 48h later.

#### Session 2

Participants came back after two days to perform the second part of the experiment. Nineteen out of the 24 came back exactly 48h later. For the remaining five, this was not possible due to personal or scanner booking contraints. Three of them came back on the same part of the day (e.g. tested in the morning on both sessions or in the afternoon on both sessions). The remaining two were tested in the morning on Session 1 but tested in the afternoon after two days. On Session 2 subjects were first reminded of the practice phase instructions, and then given instructions for the localizer phase. Both phases were performed inside the scanner.

##### Practice Phase

The practice phase was identical to the one performed in Session 1. Participants that ended Session 1 with a retrieval cycle started with a retrieval cycle, and those that ended with restudy started with restudy.

##### Functional Localizer

After the practice phase, participants were presented with all the object pictures again, with five repetitions per picture. Stimuli were presented at fovea on a white background for a duration of 2 sec, preceded by a black jittered fixation cross (0.5–7.5 sec). To make sure participants kept attending to the stimuli, a yes/no catch question appeared on the screen unpredictably. In order to not produce a high memory load, the question was always a simple question about the object shown on screen immediately before (e.g. “was the last object an instrument?”, “was the last object round?”). The localizer consisted of 730 trials: 640 of these were stimulus presentations (128 stimuli, repeated 5 times each) and 90 were catch questions. The localizer phase was divided into 5 continuous runs (not obvious to the participants). In each run the set of 128 stimuli and 18 questions were presented in a random order.

###### fMRI acquisition and pre-processing

Images were acquired on a 3T Philips Medical Systems Achieva, at the Birmingham University Imaging Centre (BUIC), using a 32-channel head-coil. Participants were instructed to avoid movement as much as possible, and head motion was further restricted by using foam pads inside the RF coil.

High resolution (1×1×1 mm) T1-weighted images were acquired for each participant at the beginning of each session, using an MPRAGE sequence (with TR =7.4 msec; TE = 3.5 msec; flip angle = 7 degrees, and FOV = 256×256×176mm).

Functional images were acquired parallel to the longitudinal axis of the hippocampus, with isotropic voxels of 3mm, a TR of 2 sec, a TE of 30 sec, and a flip angle of 80 degrees. Each volume was comprised of 38 slices with no spatial gap between them. Slices were acquired in descending order and the first five volumes of each run discarded to allow for magnetic field stabilization.

Pre-processing of the images was done using SPM12 (University College London, London, UK; http://www.fil.ion.ucl.ac.uk/spm/). Motion and time correction (slices corrected to the middle slice) were performed in this order. Data was then linearly detrended, using a Linear Model of Global Signal algorithm (Macey, Macey, Kumar, & Harper, 2004) to remove global session and voxel effects. Functional and anatomical images were co-registered. For univariate analyses, the data was further normalized to an MNI template, and finally, images were spatially smoothed with an 8-mm FWHM Gaussian kernel. The multivariate analyses were performed in the participants’ native space, that is, with no normalization or smoothing of the data.

###### Univariate analyses

Events of interest were modelled as stick functions and convolved with a canonical hemodynamic response function (HRF). Runs from the two sessions were included in the same model and session regressors were added to the general linear model (GLM) to account for differences between the two sessions. Likewise, motion parameters from spatial realignment were included as nuisance variables.

For each participant, beta values from the first level GLM were then extracted from two pre-defined anatomical regions of interest (ROIs; see ROI definition below). The average beta values from each ROI were statistically analysed with participants as random effects as described in the following. To test for fast-changing effects, we assesed whether univariate activity in a given ROI linearly increased or decreased within the first session by fitting a linear slope to the average beta values of the three retrieval and the three restudy cycles separately, individually per participant. The average slope was then tested against zero using a one-tailed dependent sample *t*-test, or compared between retrieval and restudy using a paired-sample *t*-test. This analysis identified neural changes that occurred across repeated retrieval or restudy cycles within the first day. To test for slow-changing effects across the the two day delay, the average activation from repetition 3 (last cycle on Session 1) and repetition 4 (Session 2) in each condition were subjected to a 2×2 (cycle × condition) repeated measures ANOVA. Specific effects predicted a priori (i.e., planned comparisons) were then tested using dependent sample *t*-tests (1-tailed). This latter analysis allowed us to identify neural activity that was not yet present at the end of the first scanning session but then slowly evolved across days.

### ROIs

The anatomical regions of interest (ROIs) for the univariate analyses were based on the existing sleep literature, which has shown that as memories become consolidated there is an increase in mPFC activation and a decrease in hippocampal one (Gais et al., 2007; Takashima et al., 2006). Based on this evidence, beta values were extracted from hippocampus and mPFC and tested for fast and slow changes (see above).

The hippocampi were manually traced on each participant’s structural image, using ITK-SNAP (Yushkevich et al., 2006; http://www.itksnap.org). The mPFC mask was built from a human atlas as implemented in WFUpickatlas3.0.5b (Maldjian, Laurienti, Kraft, & Burdette, 2003; http://fmri.wfubmc.edu/software/PickAtlas). The mask, composed of Broadmann Area 10, was then back-projected to the participants’ native space, using the inverse normalization parameters obtained from SPM during the segmentation step.

#### Multivariate analyses

Multivariate analyses were conducted using the RSA toolbox (Nili et al., 2014; http://www.mrc-cbu.cam.ac.uk/methods-and-resources/toolboxes/).

We first looked at semantic and episodic-specific pattern changes in the two ROIs (hippocampus and mPFC). These analyses yielded no significant results. Since we had no other specific *a priori* hypotheses regarding where in the brain the semantic and episodic-specific pattern changes would take place, we conducted two separate searchlight analyses. Searchlight analyses are ideally suited to search for regions in the brain where information is coded in a specific representational geometry (Kriegeskorte, Goebel, & Bandettini, 2006). In the case of our study, we were mainly interested in areas where multivariate patterns representing our initially studied stimuli changed across subsequent retrieval or restudy trials. All RSA analyses therefore compared the patterns elicited during initial study with the patterns elicited during subsequent retrieval and restudy cycles. Specifically, we looked for regions that increasingly (or decreasingly) coded for semantic categorical patterns, or for episodic (i.e., item-unique) patterns from the initial study to the subsequent practice (retrieval or restudy) trials. Again, we were interested in neural pattern changes that occurred either fast within-session, or slowly across sessions.

### Semantic Searchlight

To investigate how the semantic structure of memory representations changed from the original memory trace (at study) across consecutive retrieval and restudy repetitions, we conducted a searchlight analysis to find, for each cycle and each condition, where in the brain there was evidence for an increase in coding of semantic category.

The first step of this analysis consisted in building a model matrix that reflected the expected patterns of results if semanticization took place (Figure 3A). Stimuli were arranged according to their semantic category membership, in the same order across the rows and columns of the matrix (i.e. if the first four rows were dog-elephant-trumpet-accordion during encoding, the first four columns would be these same items in this same order during retrieval or restudy). Each pair of items was assigned a value, according to how similar we hypothesised them to be. Pairs of items belonging to the same semantic category (e.g. dog-elephant) were assigned a value of 1 (similar) and items belonging to different categories (dog-accordion) a value of −1 (dissimilar). Same item cells (dog-dog) were set to NaN in order to exclude any item-specific effects from this categorical analysis.

**Figure 3:**
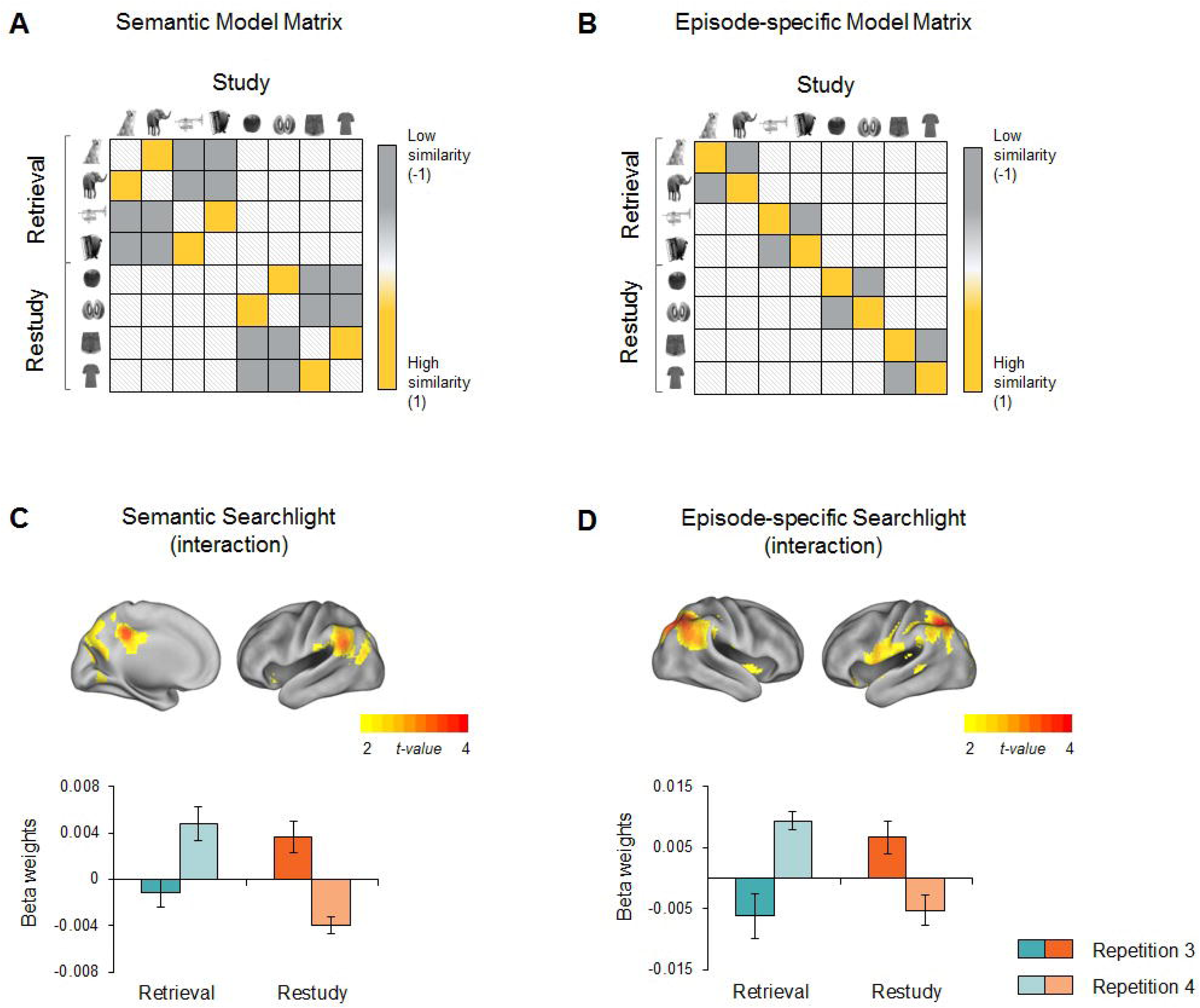
Searchlight model matrices and results for categorical information (left panel) and item-specific information (right panel). **A.** Model matrix used to investigate regions in the brain coding for categorical representations. We looked for brain areas where similarity between items belonging to the same category (yellow cells) was higher than that of items belonging to different semantic categories (filled grey cells). This matrix was correlated with the activation patterns obtained from each searchlight point to determine areas in the brain that code for categorical information in a similar fashion. **B.** Model matrix used to assess which neural regions code for item-specific information, that is, regions were similarity between the same item (yellow cells) was higher than similarity between items belonging to the same semantic category (filled grey cells). **C.** Interaction analysis ([Retrieval3 > Retrieval4] > [Restudy3 > Restudy4]) of categorical effects. We found left parietal regions (upper figure) to increasingly code for category structure during retrieval but not restudy. The lower figure shows beta weights extracted from the regions depicted above. Note that this graph has no statistical value and is used for visualisation purposes only, as to assess what drives the interaction between the two conditions. **D.** Interaction analysis of item-specific effects. Bilateral parietal regions were found to increasingly code for item unique information (upper figure), showing a strengthening of the original study trace. This pattern was found for retrieved but not restudied items (lower figure: beta weights extracted from the regions depicted above).

The searchlight ran in each participant was a sphere with a 9mm radius, collecting the multivariate patterns of activity at each voxel within the sphere. A similarity matrix was computed for each sphere using simple Pearson correlations as a metric. The matrix at each point was then correlated with the previously defined model matrix (representing the hypothesis for the categorical searchlight; Figure 3A), to determine what regions in the brain behaved similarly to the model. The resulting correlation was assigned to the centre voxel in each given sphere. As a first step, a separate searchlight was run for each retrieval and restudy cycle. From these searchlight analyses, an activation map (*r*-map) was obtained for each subject and each retrieval or restudy cycle, depicting the degree to which a given brain area showed a representational geometry (Kriegeskorte, Mur, & Bandettini, 2008) that was similar to that of the model matrix. The maps were normalized to MNI space (using the parameters from the segmentation of the T1-weighted image), smoothed with an 8mm FWHM Gaussian kernel, and statistically compared in a random-effects analysis (see next paragraph). This first analysis step resulted in 8 searchlight maps per participant, 4 representing categorical patterns during each retrieval cycle, and 4 representing categorical pattern similarity during each restudy cycle.

As a second step, group analyses were conducted on the normalized and smoothed searchlight maps from each subject within a second-level GLM. Similarly to the univariate ROI analyses, one contrast was established to assess fast-changing (within session) effects, and another one to assess slow-changing effects between the two sessions. For the within session effects we looked for the interaction of regions that increased linearly across retrieval but not restudy repetitions. For between session effects we contrasted regions that increased from repetition 3 (last cycle on day 1) to repetition 4 (day 2) in the retrieval but not the restudy condition. The results are reported at an uncorrected p-level of *p*<0.001, with a minimum extent threshold of *k*=10 voxels. Our main hypotheses all concerned similarity that were significantly more pronounced in the retrieval compared to the restudy condition. For reasons of completeness, corresponding contrasts were also created for regions showing an increase in similarity for restudy but not retrieval. These contrasts are not reported since they yielded no significant results in any of the group-level comparisons.

### Episode-Specific Searchlight

Episode-specific effects were assessed in a similar way to categorical ones. The major difference was the definition of the model matrix. In this case, we were interested in areas in the brain that coded the similarity between each item’s unique representation at study, and the same item’s subsequent retrieval or restudy representation. Accordingly, same item cells (dog-dog) were set to 1 (high similarity) whereas cells of items belonging to the same category (dog-elephant) were set to −1 (low similarity). Between category cells (dog-accordion) were set to NaN (Figure 3B).

The rest of the analysis followed the same procedure as the semantic searchlight. Results are reported at an uncorrected p-level of *p*<0.001, with a minimum extent threshold of *k*=10 voxels.

## Results

### Behavioural performance during practice

Behavioural analyses were conducted to assess performance during practice cycles. We first analysed the choice of the correct semantic category after the presentation of each stimulus pair in retrieval or restudy cycles (Table 1). A 2×4 (condition × cycle) repeated measures ANOVA was conducted and yielded a significant main effect of condition [F(1,19) = 31.56, p=.000, η^2^_partial_ =.63], a significant main effect of cycle [F(3,57) = 18.42, p=.000,η^2^_partial_ =.49] and a significant condition × cycle interaction [F(3,57) = 11.83, p=.000, η^2^_partial_ =.38]. Not surprisingly, the proportion of correct responses for restudy trials was higher than for retrieval trials (M_restudy_ = 0.89, SD= 0.08; M_retrieval_= 0.79, SD=0.13). The main effect of cycle reflects the fact that, regardless of condition, performance on the first cycle was significantly lower than performance in all subsequent ones. No other significant differences were found across cycles. The interaction shows that performance during restudy cycles remained relatively constant, while retrieval performance linearly increased from one cycle to the other across the first three repetitions and decreased again on the fourth (after two days; see Table 1).

**Table 1:** Behavioural results during the practice phase. Restudy 1 was significantly different from 2 [t(21) = −2.73, p=.012)] and 4 [t(20) = −4.32, p=.000)]. No other significant differences were found in restudy. Retrieval 1 was significantly different from all of the others (Ret1-Ret2 t(21) = −7.70, *p* =.000; Ret1-Ret3 t(20) = −7.13, *p*=.000; Ret1-Ret4 t(20) = −3.72 *p* = .001). Retrieval 2 differed from 3 (t(20) = −2.26, *p* =.035) and 3 from 4(t(19) = −2.87, *p* =.010).

|  | Overall | Restudy | Retrieval |
| --- | --- | --- | --- |
| *Correct Semantic Category* | *M (SD)* | *M (SD)* | *M (SD)* |
| Cycle 1 | 0.79 (0.09) | 0.87 (0.08) | 0.70 (0.13) |
| Cycle 2 | 0.85 (0.08) | 0.90 (0.08) | 0.81 (0.12) |
| Cycle 3 | 0.87 (0.10) | 0.89 (0.10) | 0.86 (0.11) |
| Cycle 4 | 0.85 (0.10) | 0.92 (0.07) | 0.78 (0.15) |
| *Correct Trial / Correct Category* |  |  |  |
| Cycle 1 | --- | --- | 0.75 (0.22) |
| Cycle 2 | --- | --- | 0.88 (0.10) |
| Cycle 3 | --- | --- | 0.86 (0.15) |
| Cycle 4 | --- | --- | 0.87 (0.10) |

To ensure participants were not only retrieving object category, but actually thinking back to the item associated with the retrieval cue, we measured the proportion of trials where after chosing the correct semantic category participants also reported (during the massed feedback phase) that they had thought of the correct item. In this analysis, we found a main effect of cycle [F(3,42) = 6.00, p=.002, η^2^_partial_ =.30]. Performance increased from the first retrieval attempt to subsequent ones (Table 1).

### Univariate changes in mPFC and the hippocampus

Regarding univariate effects, we hypothesised that if retrieval acts as a fast online consolidation-like mechanism, neural changes specific to this condition should occur within the first session of the experiment, and should parallel the effects reported in the sleep-dependent consolidation literature (Gais et al., 2007; Takashima et al., 2006). To test this, we computed the linear slopes of mPFC and hippocampal activation for retrieval and restudy across the first three practice cycles (Session 1 of the experiment; see Figure 1A), and tested whether retrieval slopes show a steeper increase (mPFC) or decrease (hippocampus) across retrievals compared to restudy repetitions. To test for slower changes that occur across sessions, we compared activations on the last cycle of Session 1 (repetition 3) with the practice trials in Session 2 (repetition 4). The specific predictions for each ROI, according to our fast consolidation account, are depicted in Figure 1C. We hypothesised that mPFC activation should increase during retrieval, but not restudy trials, as a reflection of retrieval’s role in embedding memories into pre-existing cortical knowledge, creating a new neocortical trace. Across sessions, the difference in mPFC activation between third and fourth repetition was expected to be greater for restudied items, given previous research showing that restudy benefits more from study than retrieval (Bäuml, Holterman, & Abel, 2014). As for hippocampal activation, if memories gradually become independent from the hippocampus, a decrease in hippocampal activation was to be expected. This decrease should be retrieval-specific and unfold within the first session if retrieval acts as a fast, online consolidation-like mechanisms. We also expected to find a retrieval-specific decrease across sessions if participants rely more on a neocortical trace and less on a hippocampal one during delayed retrieval on Session 2.

#### Medial Prefrontal Cortex

In mPFC, the retrieval slope significantly differed from zero (t(19) =3.66, p=.001), showing a within-session increase from first to third repetition. The restudy slope did not significantly differ from zero (t(19) = -.65, p=.26). The linear slopes of retrieval and restudy also differed significantly from each other (t(19) = 2.60, p=.009). The within-session univariate results thus suggest a rapid change in activity that is specific to retrieval (Figure 2A).

**Figure 2:**
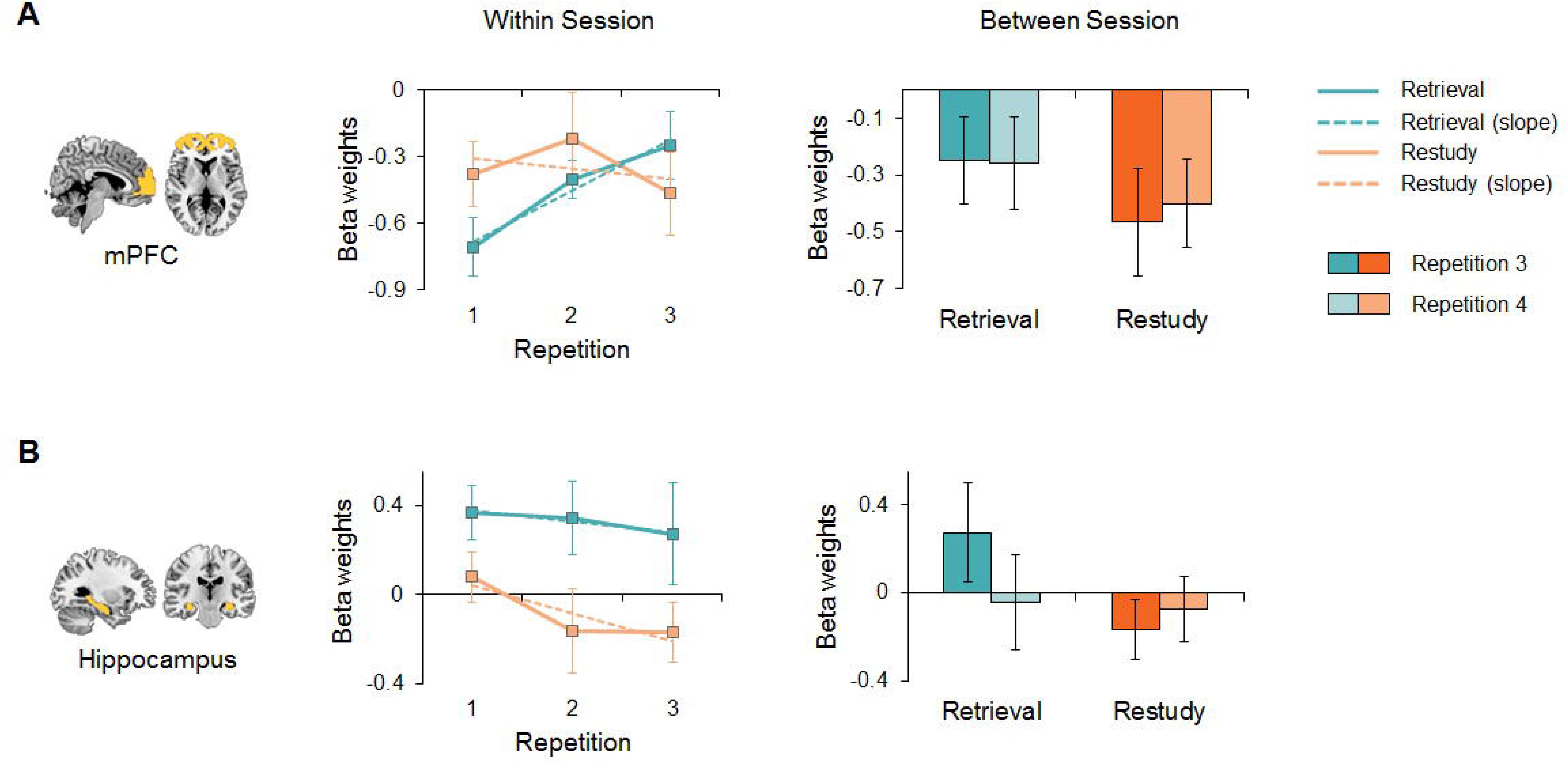
Univariate results within (left) and across (right) sessions. Retrieval results are presented in blue and restudy in orange, with the third repetition in darker colours than the fourth. **A.** mPFC results were congruent with our predictions, with mPFC activation increasing within the first session of the experiment for retrieval trials. The difference between retrieval and restudy slopes (dashed line) was statistically significant. No significant differences were found for either condition across sessions. **B.** Contrary to our predictions, hippocampus activation did not decrease within the first session, and there was no difference between the slopes of both conditions. A significant difference was found, however, between the third and fourth retrieval repetitions, with hippocampus activation decreasing across the two days. No such difference was found for restudy trials, with a marginally significant session × condition interaction.

No evidence for slowly evolving changes was found, with no significant effects across sessions in mPFC (main effect of condition: F(1,19)=1.58, p=.224; main effect of session and condition × session interaction both F<1, n.s).

#### Hippocampus

In the hippocampus, the retrieval slope was significantly different from zero (t(19) = −2.39, p=.01), whereas the restudy one was not (t = -.70, p=.25). We found no significant differences between conditions within the first session (t(19) = −1.69, p=.107), contrary to our predictions.

We did, however, find such a decrease across sessions when comparing the third practice trial (end of day 1) to the fourth practice trial (day 2). The decrease was significant for retrieved items (t(19)=2.02, p=.029; Figure 2B) but not for restudied items (t(19)=-.374, p=.357). This resulted in a marginally significant condition × cycle interaction (F(1,19)=3.48, p=.078).

### Multivariate effects

To investigate what regions in the brain increasingly coded for semantic and episode-specific effects, we ran two independent whole-brain searchlights. These searchlights compared the representational geometry at each searchlight location with a conceptual model matrix formalizing our hypotheses regarding semantic and episodic-level coding (Figures 3A and B), and specifically the level of similarity between representations at study and at each subsequent cycle of practice (retrieval or restudy). These analyses resulted in 8 activation maps (*r*-maps) per participant, representing regions in the brain that behave similarly to the model matrix during each individual retrieval and restudy cycle. These *r*-maps were then subjected to second-level group analyses, following a similar logic as for the univariate analyses. For within-session effects, we computed a linear contrast emcompassing the first three repetitions of retrieval or restudy (Session 1). To assess between-session effects, we contrasted regions where similarity with the model increased from the third to the fourth cycle (i.e., across days) for retrieval but not restudy, paralleling the univariate analyses reported above.

#### Semantic searchlight

Our fast consolidation account predicts that retrieval should lead to an increase in semantic structure, reflecting the *semanticization* of items, across consecutive retrieval cycles. If such semanticization occurs rapidly, as predicted by the fast consolidation account (Antony et al., 2017), it should be observed in neural pattern changes within Session 1 (linear contrast).

Comparing our semantic category model matrix (Figure 3A) to the empirical similarity matrices acquired from each point of the searchlight, we found no significant changes in semantic structure within session. We did, however, find that across sessions, parietal regions (Figure 3C) increasingly coded for semantic category, including clusters in bilateral cingulate gyrus, precuneus, left superior temporal gyrus, and supramarginal gyrus (Table 2).

**Table 2:** Searchlight results.

|  | XYZ | t-value | k |
| --- | --- | --- | --- |
| *Semantic Effects* |  |  |  |
|  | 3 -40 41 | 3.95 | 117 |
|  | -60 -58 26 | 3.76 | 79 |
|  | 33 -67 -25 | 3.59 | 23 |
| *Episode-specific Effects* |  |  |  |
|  | 30 -58 41 | 4.37 | 522 |
|  | -33 -58 44 | 4.32 | 289 |
|  | -21 -25 11 | 3.54 | 27 |
|  | -21 -79 14 | 3.38 | 23 |
|  | 48 -37 35 | 3.29 | 13 |

#### Episode-Specific searchlight

For episode-specific effects, the fast consolidation account predicts that retrieval leads to a differentiation of items (increased similarity with their unique study patterns compared with items from the same semantic category) across cycles, within the first session. No regions were found to increasingly code for episode-specific information within Session 1. However, strong item-unique effects were found across sessions, with mostly parietal regions encoding item identity significantly more on cycle 4 (day 2) compared to cycle 3 (day 1) of retrieval, with no difference between restudy cycles (Figure 3D). Regions implicated in episode-and retrieval-specific pattern changes included bilateral superior and inferior parietal lobe, as well as bilateral precuneus, right angular gyrus, left supramarginal and middle occipital gyrus (Table 2).

Note that some existing studies in the offline consolidation literature have also found that sleep leads to a decontextualization of memories (Cairney et al., 2011). Such decontextualization would predict a decrease in similarity with an episode’s specific study pattern. We thus also tested for retrieval-specific decreases in similarity, but found no significant changes either within-or across sessions.

## Discussion

The standard consolidation model (McClelland et al., 1995) assumes that neocortex and hippocampus interact during offline periods to stabilize and consolidate memories. Supporting this theory, sleep studies found that offline periods after learning decrease hippocampal involvement but increase medial prefrontal cortex engagement during retrieval (Gais et al., 2007; Takashima et al., 2006). These findings suggest that hippocampal-neocortical replay during sleep leads to the creation of a new neocortical trace, rendering memories less hippocampus-dependent (Frankland & Bontempi, 2005). The present study is the first empirical test of a recent framework, proposing that retrieval exerts its memory benefits through replay mechanisms similar to those involved in offline consolidation, acting as a fast route to memory consolidation (Antony et al., 2017).

First we tested whether univariate changes in brain activity, induced by repeated retrival, mirrored results reported in the sleep literature (e.g. Gais et al., 2007). We found an increase in mPFC activation across successive retrieval attempts within the first session, while no increase was found for restudy. This observation is consistent with the hypothesis that retrieval engages consolidation-like mechanisms, leading to the rapid creation of a neocortical trace that can be ultilized for future remote access to the memory. Additionally, we found a decrease in hippocampal activation across, rather than within, sessions. This finding does not lend direct support to the fast consolidation account, suggesting that at short delays, active retrieval still relies on the hippocampal trace. Retrieval did become less hippocampus-dependent but in a slowly evolving fashion, evident only after a two-day delay.

These univariate changes cannot be easily explained by effort or difficulty based interpretations. Such interpretations would predict an increase rather than decrease in hippocampal engagement after a delay of several days, since retrieval is rendered more difficult then (as evidenced by the drop in behavioural performance on day 2). Similarly, it could be argued that the retrieval-specific increase in mPFC activation is simply reflecting retrieval being initially more demanding than restudy, but becoming easier across repetitions, associated with an increase in default mode regions (Fox et al., 2005). Contrarily, we found that mPFC activity remained elevated across the two-days delay, when the task is again rendered more difficult.

Taken together, our univariate results are consistent with an online role of retrieval in supporting memory consolidation. The slow hippocampal decrease indicates that the benefits of this stabilization for episodic retrieval only become relevant at longer delays, when the hippocampal trace is presumably less accessible. At shorter delays, the neocortical and hippocampal traces seem to co-exist (as previously suggested by Winocur, Moscovitch and Bontempi (2010)).

On the level of distributed representations, we were interested in qualitative changes in representational geometries, induced by repeated retrieval at faster and slower timescales. Specifically, we tested whether retrieval opens an opportunity for items to become embedded into pre-existing neocortical knowledge structures, as shown in the sleep literature (Schapiro et al., 2017). If so, similarity between items belonging to the same semantic category (e.g. animals) should increase. Although we expected these changes to occur online (i.e., across retrievals within Session 1), we found them exclusively across sessions. The increase in categorical coding across retrieval trials was most pronounced in parietal regions, in the core recollection network (Rugg & Vilberg, 2012), consistent with these regions’ role in vivid recollection (Kuhl & Chun, 2014), particularly after long delays (Brodt et al., 2016; Oedekoven, Keidel, Berens, & Bird, 2017).

A similar pattern emerged regarding our third question of whether episode-unique information would be strengthened by differentiation, as predicted by our fast consolidation account (Antony et al., 2017), or gradually lost, as suggested by some sleep studies (Cairney et al., 2011; but see Jurewicz et al., 2016). Corrobortating our prediction, idiosyncratic study traces were strengthened throughout subsequent retrievals in superior parietal regions, related to familiarity-based remembering (Hutchinson, Uncapher, & Wagner, 2009) and the subjective perception of remembering (Wagner, Shannon, Kahn, Buckner, & Louis, 2005). This pattern change was again only found across a two-days delay, indicating a slowly evolving change in the underlying memory trace, and potentially in the quality of remembering.

Our pattern fMRI results indicate that retrieval concurrently strengthens semantic-categorical and episodic aspects of memories. This is consistent with recent findings in the offline literature, showing that episodic and semantic effects are not mutually exclusive but can co-occur (Schlichting, Mumford, & Preston, 2015; Schapiro et al., 2017; Tompary & Davachi, 2017). In an fMRI sleep study, Schapiro and colleagues (2017) showed that sleep enhances categorical structure, at no expense of the item-unique representation, which is preserved after sleep. Similarly, Tompary and Davachi (2017) found, after a week, both the reinstatement of item-specific encoding patterns and a similarity increase between patterns evoked during the retrieval of overlapping *(vs.* non-overlapping) memories. Together with our findings, evidence supporting the co-existence of two cortical traces accumulates, consistent with a multiple trace view (Winocur et al., 2010).

Our multi-day design allowed to explicitly investigate the timescale at which retrieval induces neural pattern changes. Surprisingly, all representational changes, as well as the univariate hippocampal disengagement, occurred slowly across days, suggesting that retrieval exerts its longterm benefits by interacting with sleep-or time-dependent processes. For example, it is possible that repeatedly retrieved memories are “tagged” for prioritized replay during subsequent sleep, leading to more stabilisation than for non-retrieved or restudied memories. While such a tagging account can explain why retrieval’s benefits are typically only found after long delays, empirical findings have shown that retrieved memories actually benefit less from subsequent sleep than restudied ones (Bäuml et al., 2014). A tagging account can thus not fully explain existing results in the literature. Given the many parallels between retrieval and offline consolidation (Antony et al., 2017), we favour the view that memory reactivation during retrieval changes the underlying memory in a way similar to offline replay. Replay could induce plasticity immediately, but the resulting neural changes might only become relevant and visible at a later time point, when access to the memory trace is more difficult and relies relatively more on neocortical access routes.

To our knowledge, this study is the first direct test of the fast consolidation account of retrieval-mediated learning, contrasting differential effects of retrieval and restudy at fast and slow timescales. Previous fMRI studies have investigated the effects of repeated retrieval practice on univariate (Wing, Marsh, & Cabeza, 2013) and multivariate (Jonker, Dimsdale-Zucker, Ritchey, Clarke, & Ranganath, 2018.; Lee, Samide, Richter, & Kuhl, 2017) activity patterns. Many of their results are consistent with the present study. Wing et al. (2013) showed stronger connectivity between hippocampus and mPFC after retrieval than relearning. Although this finding was not interpreted within a consolidation framework, it parallels results in the offline consolidation literature (Gais et al., 2007). Lee and colleagues (2017) investigated how multivatiate pattern changes at item and categorical levels across repeated retrievals predicted later memory enhancement, compared to no practice. Neural reactivation of item-specific information was related to a behavioural increase in correct rejections, while reactivation of categorical patterns related to an increase in false alarms to similar lures. Jonker et al. (2018) compared pattern changes across retrieval and restudy opportunities, and found that retrieval strengthens the neural representations of the target objects along with contextually (but not semantically) linked objects in parietal cortex. In both studies retrieval-induced changes were most pronounced in parietal cortex. Together with our searchlight results, these findings suggest that retrieval shapes representations by extracting commonalities between new memories and previously stored knowledge, while preserving relevant stimulus-specific information. They also highlight the role of parietal cortex as a hub for storing and combining different types of information.

We belive our work is an important step towards a neurobiologically plausible mechanism for retrieval-mediated strengthening, and can inform cognitive theories of the testing-effect (Karpicke et al., 2017). Similar to the semantic mediator and the elaborative retrieval accounts (Carpenter, 2009), the rapid consolidation framework assumes that retrieval facilitates the integration of new memories with existing neocortical knowledge structures by co-activating related information. On the other hand, our categorical searchlight results are in line with a contextual reinstatement account (Rowland, 2014) by showing a retrieval-specific increase in episodic reinstatement. Our work may help unifying these accounts, demonstrating that retrieval can affect a memory representation at multiple levels, from global-categorical to idiosyncratic episode-specific features.

In sum, we empirically tested the fast consolidation account of retrieval. We show that retrieval leads to a rapid increase in mPFC activation, consistent with slow changes described in the sleep literature (Gais et al., 2007; Takashima et al., 2006). In line with this literature (Frankland & Bontempi, 2005), we argue that retrieval aids the creation of a neocortical trace which, over longer periods of time, offers an additional access route, rendering memories less hippocampus-dependent, as evidenced by the decrease in hippocampal activation across sessions. We also show that retrieval strengthens both semantic and episode-specific neural patterns, suggesting these traces can co-exist and might play a more or less important role at short and long delays. Our findings are congruent with a consolidation-like mechanism, although not necessarily a fast acting one.

## Aknowledgments

This work was supported by grant ES/M001644/1, from the Economical and Social Research Council UK, awarded to M.W.

## References

Antony, J. W., Ferreira, C. S., Norman, K. A., & Wimber, M. (2017). Retrieval as a Fast Route to Memory Consolidation. Trends in Cognitive Sciences, 21(8), 573–576.

Bäuml, K.-H., Holterman, C., & Abel, M. (2014). Sleep can reduce the testing effect: It enhances recall of restudied items but can leave recall of retrieved items unaffected. Journal of Experimental Psychology: Learning, Memory and Cognition, 40(6), 1568–1581.

Brodeur, M. B., Guérard, K., & Bouras, M. (2014). Bank of Standardized Stimuli (BOSS) Phase II: 930 New Normative Photos. PLoS ONE, 9(9), e106953.

Brodt, S., Pöhlchen, D., Flanagin, V. L., Glasauer, S., Gais, S., & Schönauer, M. (2016). Rapid and independent memory formation in the parietal cortex. Proceedings of the National Academy of Sciences, 113(46), 13251–13256.

Cairney, S. A., Durrant, S. J., Musgrove, H., & Lewis, P. A. (2011). Sleep and environmental context: Interactive effects for memory. Experimental Brain Research, 214(1), 83–92.

Carpenter, S. K. (2009). Cue strength as a moderator of the testing effect: The benefits of elaborative retrieval. Journal of Experimental Psychology: Learning, Memory, and Cognition, 35(6), 1563–1569.

Dudai, Y., Karni, A., & Born, J. (2015). The Consolidation and Transformation of Memory. Neuron, 88(1), 20–32.

Fox, M. D., Snyder, A. Z., Vincent, J. L., Corbetta, M., Van Essen, D. C., & Raichle, M. E. (2005). The human brain is intrinsically organized into dynamic, anticorrelated functional networks. Proceedings of the National Academy of Sciences, 102(27), 9673–9678.

Frankland, P. W., & Bontempi, B. (2005). The organization of recent and remote memories. Nature Reviews Neuroscience, 6(2), 119–130.

Gais, S., Albouy, G., Boly, M., Dang-Vu, T. T., Darsaud, A., Desseilles, M., … Peigneux, P. (2007). Sleep transforms the cerebral trace of declarative memories. Proceedings of the National Academy of Sciences, 104(47), 18778–18783.

Hulbert, J. C., & Norman, K. A. (2015). Neural Differentiation Tracks Improved Recall of Competing Memories Following Interleaved Study and Retrieval Practice. Cerebral Cortex, 25, 3994–4008.

Hutchinson, J. B., Uncapher, M. R., & Wagner, A. D. (2009). Posterior parietal cortex and episodic retrieval: Convergent and divergent effects of attention and memory. Learning and Memory, 16(6), 343–356.

Jonker, T. R., Dimsdale-Zucker, H., Ritchey, M., Clarke, A., & Ranganath, C. (2018). Neural reactivation in parietal cortex enhances memory for episodically linked information. Proceedings of the National Academy of Sciences, 115(43), 11084–11089.

Jurewicz, K., Cordi, M. J., Staudigl, T., & Rasch, B. (2016). No Evidence for Memory Decontextualization across One Night of Sleep. Frontiers in Human Neuroscience, 10, 7.

Karpicke, J. D., Lafayette, W., & States, U. (2017). Retrieval-Based Learning: A Decade of Progress. Learning and Memory: A Comprehensive Reference (Third Edit). Elsevier.

Kriegeskorte, N., Goebel, R., & Bandettini, P. (2006). Information-based functional brain mapping. Proceedings of the National Academy of Sciences, 103(10), 3863–3868.

Kriegeskorte, N., Mur, M., & Bandettini, P. (2008). Representational similarity analysis – connecting the branches of systems neuroscience. Frontiers in Systems Neuroscience, 2(4), 1–28.

Kuhl, B. A., & Chun, M. M. (2014). Successful Remembering Elicits Event-Specific Activity Patterns in Lateral Parietal Cortex. Journal of Neuroscience, 34(23), 8051–8060.

Lee, H., Samide, R., Richter, F. R., & Kuhl, B. A. (2017). Decomposing parietal memory reactivation to predict consequences of remembering. Cerebral Cortex, 1–14.

Macey, P. M., Macey, K. E., Kumar, R., & Harper, R. M. (2004). A method for removal of global effects from fMRI time series. NeuroImage, 22(1), 360–366.

Maldjian, J. A., Laurienti, P. J., Kraft, R. A., & Burdette, J. H. (2003). An automated method for neuroanatomic and cytoarchitectonic atlas-based interrogation of fMRI data sets. Neuroimage, 19(3), 1233–1239.

McClelland, J. L., McNaughton, B. L., & O’Reilly, R. C. (1995). Why there are complementary learning systems in the hippocampus and neocortex: Insights from the successes and failures of connectionist models of learning and memory. Psychological Review, 102(3), 419–457.

Nieuwenhuis, I. L. C., & Takashima, A. (2011). The role of the ventromedial prefrontal cortex in memory consolidation. Behavioural Brain Research, 218(2), 325–334.

Nili, H., Wingfield, C., Walther, A., Su, L., Marslen-Wilson, W., & Kriegeskorte, N. (2014). A Toolbox for Representational Similarity Analysis. PLoS Computational Biology, 10(4).

Oedekoven, C. S. H., Keidel, J. L., Berens, S. C., & Bird, C. M. (2017). Reinstatement of memory representations for lifelike events over the course of a week. Scientific Reports, 7, 14305.

Peirce, J. W. (2006). PsychoPy—Psychophysics software in Python. Journal of Neuroscience Methods, 162, 8–13.

Rasch, B., Büchel, C., Gais, S., & Born, J. (2007). Odor Cues During Slow-Wave Sleep Prompt Declarative Memory Consolidation. Science, 315(5817), 1423–1426.

Roediger, H. L., & Karpicke, J. D. (2006). The Power of Testing Memory Basic Research and Implications for Educational Practice. Perspectives in Psychological Science, 1(3), 181–210.

Rowland, C. A., & Rowland, C. A. (2014). Psychological Bulletin The Effect of Testing Versus Restudy on Retention: A Meta-Analytic Review of the Testing Effect The Effect of Testing Versus Restudy on Retention: A Meta-Analytic Review of the Testing Effect, 140(6), 1432–1463.

Rugg, M. D., & Vilberg, K. L. (2013). Brain Networks Underlying Episodic Memory Retrieval. Current Opinion in Neurobiology, 23(2), 255–260.

Schapiro, A. C., McDevitt, E. A., Chen, L., Norman, K. A., Mednick, S. C., & Rogers, T. T. (2017). Sleep Benefits Memory for Semantic Category Structure while Preserving Exemplar-Specific Information. Scientific Reports, 7(1), 1–22.

Schlichting, M. L., Mumford, J. A., & Preston, A. R. (2015). Learning-related representational changes reveal dissociable integration and separation signatures in the hippocampus and prefrontal cortex. Nature Communications, 6, 1–10.

Takashima, A., Petersson, K. M., Rutters, F., Tendolkar, I., Jensen, O., Zwarts, M. J., … Fernández, G. (2006). Declarative memory consolidation in humans: A prospective functional magnetic resonance imaging study. Proceedings of the National Academy of Sciences, 103(3), 756–761.

Tompary, A., & Davachi, L. (2017). Consolidation Promotes the Emergence of Representational Overlap in the Hippocampus and Medial Prefrontal Cortex. Neuron, 96(1), 228–241.

Wagner, A. D., Shannon, B. J., Kahn, I., Buckner, R. L., & Louis, S. (2005). Parietal lobe contributions to episodic memory retrieval. Trends in Cognitive Science, 9(9), 445–453.

Wing, E. A., Marsh, E. J., & Cabeza, R. (2013). Neural correlates of retrieval-based memory enhancement: An fMRI study of the testing effect. Neuropsychologia, 51(12), 2360–2370.

Winocur, G., Moscovitch, M., & Bontempi, B. (2010). Memory formation and long-term retention in humans and animals: Convergence towards a transformation account of hippocampal-neocortical interactions. Neuropsychologia, 48(8), 2339–2356.

Xiao, J., Hays, J., Ehinger, K., Oliva, A., & Torralba, A. (2010). SUN Database: Large-scale Scene Recognition from Abbey to Zoo. In IEEE Conference on Computer Vision and Pattern Recognition.

Yushkevich, P. A., Piven, J., Hazlett, H. C., Smith, R. G., Ho, S., Gee, J. C., & Gerig, G. (2006). User-guided 3D active contour segmentation of anatomical structures: Significantly improved efficiency and reliability. Neuroimage,31(3), 1116–1128.

